# An analysis of RNA quality metrics in human brain tissue

**DOI:** 10.1101/2024.10.14.618253

**Authors:** Jiahe Tian, Tiffany G. Lam, Sophie K. Ross, Benjamin Ciener, Sandra Leskinen, Sharanya Sivakumar, David A. Bennett, Vilas Menon, Guy M. McKhann, Alexi Runnels, Andrew F. Teich

**Author notes:** Corresponding author Andrew F. Teich.

## Abstract

Human brain tissue studies have historically used a range of metrics to assess RNA quality. However, few large-scale cross-comparisons of pre-sequencing quality metrics with RNA-seq quality have been published. Here, we analyze how well metrics gathered before RNA sequencing (post-mortem interval (PMI) and RNA integrity number RIN) relate to analyses of RNA quality after sequencing (Percent of counts in Top Ten genes (PTT), 5’ bias, and 3’ bias) as well as with individual gene counts across the transcriptome. We conduct this analysis across four different human cortical brain tissue collections sequenced with varying library preparation protocols. PMI and RIN have a low inverse correlation, and both PMI and RIN show consistent and opposing correlations with PTT. Unlike PMI, RIN shows strong consistent correlations with measurements of 3’ and 5’ bias, and RIN also correlates with 3,933 genes across datasets, in comparison to 138 genes for PMI. Neuronal and immune response genes correlate positively and negatively with RIN respectively, suggesting that different gene sets have divergent relationships with RIN in brain tissue. In summary, these analyses suggest that conventional metrics of RNA quality have varying degrees of value, and that PMI has an overall minimal but reproducible effect on RNA quality.

## Introduction

Brain tissue is routinely collected and stored at academic hospitals for a variety of research programs (1, 2). A major determinant of tissue quality is the post-mortem interval (PMI), which is often assumed to be relevant for RNA quality. For RNA studies, an RNA integrity number (RIN), as measured by a bioanalyzer, is frequently used as a more direct metric.

The relationship between RIN and PMI on the one hand and RNA quality on the other has been extensively studied by multiple groups. While RIN has been shown to consistently predict RNA quality (3-6), the literature on PMI is more mixed. Several groups have found little to no relationship between PMI and RNA quality (4, 6-9), while others have found that PMI does affect the expression of some genes (3, 10). Notably, comparison of RIN and PMI to RNA quality is mostly limited to qPCR studies (3, 5-9, 11, 12), and some microarray data (7, 13). With RNA-seq now ubiquitous as a technology (14-17), there is a need for a more thorough comparison of these pre-sequencing quality metrics with commonly used QC metrics relevant for sequencing-based transcriptomic analysis in brain tissue, as well as an analysis of how RIN and PMI relate to gene expression across the entire transcriptome.

Here, we analyze how well PMI and RIN relate to analyses of RNA quality in cortical brain tissue after RNA-seq. We do this across four different datasets of RNA-seq data, including 106 surgical specimens with poly-A selection, 534 autopsy specimens with poly-A selection, and two different autopsy datasets with ribosomal RNA depletion and no poly-A selection, with 513 and 365 specimens in each dataset. We relate PMI and RIN to a range of post-sequencing QC metrics and show that RIN is predictive of overall RNA quality, while PMI has a reproducible but overall minimal impact. We also identify genes that correlate most reliably with RIN across all four datasets, which has implications for research where these genes are relevant. In summary, our work is the most extensive analysis to date of the effects of RIN and PMI in human brain tissue and has implications for the relative importance that brain banks should place on these parameters.

## Materials and Methods

*Data Acquisition:* All four datasets used in this study were generated for other studies. The NPH dataset is a collection of cortical biopsies taken during ventricular shunt placement at Columbia University Irving Medical Center. Biopsies were taken from frontal cortex in 2/3 of the subjects and parietal cortex in 1/3 of the subjects in our original cohort. As reported in (16), RNA was extracted from biopsy samples using miRNeasy Mini Kit (QIAGEN; Cat No./ID: 217004), and samples with RIN values ≥ 6 were selected for sequencing. RNAs were prepared for sequencing using the Illumina TruSeq mRNA library prep kit, and samples underwent single-end sequencing to 30M read depth.

New York Brain Bank (NYBB) RNA-seq data from dorsolateral/dorsomedial prefrontal cortex was also generated at Columbia University (18). Total RNA was used to make cDNA using Kapa Total library prep, which removes ribosomal RNA and is optimized for degraded RNA. Our QC threshold for using RNA was based on RIN, trace degradation, and presence of ribosomal RNA subunits (some samples had a RQN generated on a fragment analyzer instead of RIN on a bioanalyzer, and we use “RIN” to refer to both RIN and RQN values from this dataset given the interchangeability of these two numbers). RNA samples that had RIN above 4 passed. Samples that had RIN values between 2.5-3.9 were evaluated and passed for sequencing based on visual inspection of trace degradation and ribosomal subunit presence.

Religious Orders Study and Rush Memory and Aging Project (ROSMAP) RNA-seq data from dorsolateral prefrontal cortex (DLPFC) was obtained from the Rush Alzheimer’s Disease Center (19, 20); ROSMAP RNA-seq data is available via the AMP-AD data portal through Synapse (https://www.synapse.org/#!Synapse:syn3219045) and the RADC Research Resource Sharing Hub (https://www.radc.rush.edu). Of the 899 ROSMAP samples in this study, 534 were sequenced using poly-A selection, while 365 used KAPA Stranded RNA-Seq Kit with RiboErase (kapabiosystems); these two sets are analyzed separately in this manuscript.

All FASTQ files were aligned to the GRCh38 genome (Ensembl Release 101) using STAR (v2.7.6a) in 2-pass mapping mode with standard ENCODE parameters. Gene counts were quantified from STAR-aligned BAM files using featureCounts (v2.0.1). RNA sequencing metrics were collected using Picard by generating a ribosomal interval list and a ref_flat file from annotated genomic data. The median 5’ and 3’ bias of the 1000 most highly expressed transcripts in each biopsy sample were extracted for further analysis.

### Correlation Analysis on Datasets

Correlation analysis was conducted on NPH (*n* = 106), NYBB (*n* = 513), and ROSMAP (*n* = 534, 365) datasets to examine relationships among metrics. Spearman’s correlation analysis was performed using the corr.test function from the psych library in R, with p-values adjusted for false discovery rate (FDR) at a threshold of 0.05. The metrics analyzed included RNA Integrity Number (RIN), the proportion of the top 10 gene counts to total gene expression (PTT), Post-Mortem Interval (PMI), DV200, median 5’ bias, median 3’ bias, and the median 5’ to 3’ ratio, where applicable. For Tables 1 and 2, each panel (a, b, c, etc.) has p-values FDR adjusted across that panel.

**Table 1:** Correlations of QC Metrics in NPH Biopsy Data and NYBB Autopsy Data.

(a) NPH (n = 106)
|  | 1 | 2 | 3 | 4 | 5 |
| --- | --- | --- | --- | --- | --- |
| 1. RIN | 1.00 |  |  |  |  |
| 2. PTT | -.55** | 1.00 |  |  |  |
| 3. median 5' bias | -.025 | -.54** | 1.00 |  |  |
| 4. median 3' bias | -.18 | .57** | -.75** | 1.00 |  |
| 5. median 5' to 3' bias ratio | .028 | -.58** | .98** | -.78** | 1.00 |

(b) NYBB (n = 513)
|  | 1 | 2 | 3 | 4 |
| --- | --- | --- | --- | --- |
| 1. RIN | 1.00 |  |  |  |
| 2. PMI | -.11* | 1.00 |  |  |
| 3. PTT | -.53** | .19** | 1.00 |  |
| 4. DV200 | .72** | -.19** | -.60** | 1.00 |

(c) NYBB (RIN ≥ 6, n = 160)
|  | 1 | 2 | 3 | 4 |
| --- | --- | --- | --- | --- |
| 1. RIN | 1.00 |  |  |  |
| 2. PMI | -.10 | 1.00 |  |  |
| 3. PTT | -.35** | .21* | 1.00 |  |
| 4. DV200 | .58** | -.11 | -.41** | 1.00 |

\* *FDR-adjusted p-value* < 0.05, \*\* *FDR-adjusted p-value* < 0.001

**Table 2:** Correlations of QC Metrics in ROSMAP Autopsy Data.

(a) ROSMAP poly-A Selection Library Preparation (n = 534)
|  | 1 | 2 | 3 | 4 | 5 | 6 |
| --- | --- | --- | --- | --- | --- | --- |
| 1. RIN | 1.00 |  |  |  |  |  |
| 2. PMI | -.26** | 1.00 |  |  |  |  |
| 3. PTT | -.68** | .22** | 1.00 |  |  |  |
| 4. median 5' bias | .47** | .051 | -.60** | 1.00 |  |  |
| 5. median 3' bias | -.029 | -.11* | .32** | -.46** | 1.00 |  |
| 6. median 5' to 3' bias ratio | .18** | .11* | -.46** | .73** | -.92** | 1.00 |

(b) ROSMAP poly-A Selection Library Preparation (RIN ≥ 6.0, n = 441)
|  | 1 | 2 | 3 | 4 | 5 | 6 |
| --- | --- | --- | --- | --- | --- | --- |
| 1. RIN | 1.00 |  |  |  |  |  |
| 2. PMI | -.26** | 1.00 |  |  |  |  |
| 3. PTT | -.67** | .24** | 1.00 |  |  |  |
| 4. median 5' bias | .49** | .024 | -.60** | 1.00 |  |  |
| 5. median 3' bias | -.20** | -.061 | .42** | -.54** | 1.00 |  |
| 6. median 5' to 3' bias ratio | .29** | .066 | -.51** | .76** | -.93** | 1.00 |

(c) ROSMAP Ribosomal Depletion Library Preparation (n = 365)
|  | 1 | 2 | 3 |
| --- | --- | --- | --- |
| 1. RIN | 1.00 |  |  |
| 2. PMI | -.088 | 1.00 |  |
| 3. PTT | -.54** | .026 | 1.00 |

(d) ROSMAP Ribosomal Depletion Library Preparation (RIN ≥ 6.0, n = 103)
|  | 1 | 2 | 3 |
| --- | --- | --- | --- |
| 1. RIN | 1.00 |  |  |
| 2. PMI | -.0050 | 1.00 |  |
| 3. PTT | -.063 | .071 | 1.00 |

\* FDR-adjusted $p$ -value $< 0.05$ , \*\* FDR-adjusted $p$ -value $< 0.001$

### Single-gene Correlation Analysis

Spearman’s correlation analysis was performed to examine the relationship between RNA Integrity Number (RIN) and Post-Mortem Interval (PMI) for NYBB and ROSMAP samples, and RIN for NPH biopsies. Genes with fewer than 5 counts in at least 90% of samples were filtered out from the raw count matrix. The filtered counts were then normalized using variance stabilizing transformation (VST) with the *varianceStabilizingTransformation* function from the DESeq2 R package. RIN and PMI for each sample were correlated with the normalized gene counts of each gene across all samples. Ontology analysis on biological processes was performed using Enrichr (21).

## Results

We analyzed bulk RNA-seq data from neocortex across four different groups: a set of 106 biopsy specimens from patients with normal pressure hydrocephalus (16), a cohort of 899 dorsolateral prefrontal cortex specimens from the ROSMAP cohort (19, 20) (divided into two groups, see below), and a group of 513 dorsolateral/dorsomedial prefrontal cortex specimens from the New York Brain Bank (NYBB) at Columbia University. The NPH and 534 of the ROSMAP specimens underwent poly-A selection and subsequent RNA-seq, while the NYBB specimens and 365 of the ROSMAP specimens underwent enzymatic depletion of ribosomal RNA and did not employ poly-A selection. All samples had RIN values generated (see Methods), and for NYBB samples a DV200 value (the percentage of RNA fragments > 200 nucleotides) was also generated. Post-mortem interval (relevant for the three autopsy cohorts) was also recorded for all samples. After bulk RNA-seq, the percent of counts in the top ten genes (PTT) was calculated for each sample. In addition, the percent of counts in the 5’ or 3’ end of each transcript normalized by counts over the entire transcript was calculated for the two datasets with poly-A selection, given that this technique is sensitive to 3’ bias with degraded RNA (22).

The goal of this study was to examine how all of these metrics relate to each other across these four different datasets and to determine how well pre-sequencing RNA quality metrics predict RNA quality measured post-sequencing. For all autopsy datasets, we performed our analysis in both the full dataset and in a subset of samples with RIN values restricted to the range in the NPH data (RIN of 6 and higher) to evaluate if lower RIN scores in the autopsy tissue is influencing any differences with the surgical tissue set.

### RIN and PMI show similar and divergent relationships with post-sequencing QC metrics

The set of 106 surgical specimens is free of post-mortem artifact by definition, and there is no associated PMI. In this dataset, RIN correlates strongly and inversely with PTT (r = -0.55) as expected, but has no significant correlation with measures of 5’ or 3’ bias (Table 1A). One caveat to this analysis is that the correlation of 3’ bias with RIN (r = -0.18) has an adjusted p-value of 0.091. The ROSMAP dataset (discussed below) has a similar correlation between RIN and 3’ bias (r = -0.2) for higher RIN samples that is significant. The ROSMAP cohort is several fold larger than the NPH sample set, so one could speculate that the lack of significance in the NPH samples is primarily a power issue. Although 5’ and 3’ have no significant relationship with RIN in the NPH data, they are relevant RNA quality metrics in this dataset, as indicated by the fact that 5’ bias, 3’ bias, and the ratio of 5’ to 3’ reads all correlate strongly and consistently with PTT.

Next, we examined the NYBB set of RNA-seq data (Table 1B). Measurements of 5’ and 3’ bias are irrelevant in this dataset given the fact that poly-A selection was not used. Thus, for post-sequencing measurements we only have PTT. However, for pre-sequencing measurements, the DV200 was also recorded for these samples. Comparing all these available metrics to one another, we observed that RIN negatively correlates with PTT (r = -0.53), similarly to the NPH dataset. RIN and DV200 are closely aligned (r = 0.72), and RIN has a weak negative correlation with PMI (r = -0.11). Of note, PMI positively correlates with PTT in this dataset (r = 0.19), suggesting that PMI does influence RNA quality post-sequencing. When we restrict NYBB samples to the range of RIN values seen in the NPH surgical tissue dataset (RIN > 6), we see similar trends, although we lose significance for some correlations, likely due to lower power (Table 1C).

Both the NPH and NYBB RNA-seq data were generated from samples at Columbia University. To further generalize our results, we next analyzed pre-and post-sequencing RNA quality metrics in ROSMAP autopsy tissue (Table 2). The ROSMAP samples sequenced with poly-A selection demonstrated consistent and significant correlations between RIN and all other measured metrics in this dataset. PMI correlates significantly with PTT (r = 0.22). The relationship of RIN and PMI to 5’ and 3’ bias is more complex. RIN shows consistent and significant correlations with these metrics, although 3’ bias isn’t significant in the full dataset but is when we restrict our analyses to samples to the range of RIN values seen in the NPH surgical tissue dataset (RIN of 6 or higher). PMI also shows a complex relationship with 3’ bias, as it shows a significant negative correlation with 3’ bias in the full dataset (the opposite of what would be predicted), and this relationship is no longer significant when samples are restricted to RIN of 6 and higher. Although it is somewhat difficult to interpret this data, both 3’ and 5’ bias do have strong and consistent relationships with PTT for both the full and restricted datasets, suggesting that these measurements of RNA quality are performing as expected. At minimum, PMI does not relate to 3’ or 5’ bias in a coherent way, while RIN has an overall consistent trend that may depend on additional factors for reproducibility.

In the ROSMAP subjects that were sequenced with ribosomal RNA depletion, RIN correlates with PTT in the full dataset, and this is lost after restricting the samples to RIN of 6 or higher, possibly due to lower statistical power. In summary, PMI negatively correlates with RIN in two of the three autopsy datasets we examined. Both RIN and PMI correlate with PTT in all but the final dataset (ROSMAP with ribosomal RNA depletion), where only RIN correlates with PTT. The major area of divergence in predictive power of RIN vs. PMI is in measurements of 3’ and 5’ bias, where RIN significantly correlates with these measurements in the autopsy cohort and approaches significance in the NPH data, whereas PMI shows no predictive power.

### RIN and PMI show divergent effects at the individual gene level

Finally, we analyzed RNA-seq data for genes that correlate with RIN or PMI. Spearman’s correlations with PMI and RIN were calculated for all genes in the transcriptome for all four datasets, and significance was set at FDR < 0.05. We then determined how many genes positively correlate with RIN in all four datasets and found 2,512 genes that commonly overlap (see Supplemental Data for genes that commonly correlate with RIN and PMI in the four datasets). Notably, 1,421 genes negatively correlate with RIN in all four datasets, suggesting a complex relationship between RIN and gene expression values. We next determined how many genes positively and negatively correlate with PMI in the three autopsy cohorts. Interestingly, far fewer genes correlate with PMI across these datasets, with 138 positively correlating genes and no negatively correlating genes passing FDR of 0.05. To understand whether there are common biological processes that characterize the sets of genes that correlate with RIN and PMI, we performed ontology analysis on the genes in common (Table 3; See Supplemental Data for full ontology analysis). Genes that positively correlate with RIN were significantly enriched for ontology categories characterized by neuronal function, whereas genes that negatively correlate with RIN are characterized by the immune response. Given that these tissue collections are from neurodegeneration studies, we asked whether this gene signature was capturing neurodegenerative change in any way. While RIN mildly correlates with Braak stage in the ROSMAP poly-A dataset (r = -0.11, p = 0.01), RIN does not correlate with Braak stage in the ROSMAP ribosomal RNA depletion dataset (r = -0.091, p = 0.15) or the NYBB dataset (r = -0.052, p = 0.318). We have previously quantified β-amyloid and tau pathology in the NPH biopsies (16), and RIN does not correlate with either of these metrics (β-amyloid r = -0.119, p = 0.386; tau r = -0.046, p = 0.808). Thus, it is unlikely that neurodegenerative disease is indirectly driving these changes in gene expression. Instead, these ontology themes suggest that inflammation may be related to RNA degradation in brain tissue more broadly (see Discussion). The more limited sets of genes that correlate with PMI did not show significant ontology categories. In summary, the relative abundance of genes that correlate with RIN vs. PMI and the correlation of RIN with measurements of 3’ and 5’ bias point to ways that RIN is uniquely capturing RNA fragmentation in brain tissue, although PMI may also relate to some aspects of RNA degradation.

**Table 3:** Ontology Categories of Commonly Correlating Genes in the NPH, NYBB, and both ROSMAP datasets.

(a) Representative Ontology Categories (RIN Positively Correlating Genes)
| **Ontology Categories** | **FDR-adjusted**<br>**p-value** | **Odds Ratio** |
| --- | --- | --- |
| Chemical Synaptic Transmission<br>(GO:0007268) | 1.55E-23 | 4.43 |
| Anterograde Trans-Synaptic<br>Signaling (GO:0098916) | 1.34E-17 | 4.50 |
| Vesicle-Mediated Transport<br>(GO:0016192) | 4.42E-08 | 2.29 |
| Neurotransmitter Secretion<br>(GO:0007269) | 6.33E-07 | 7.35 |
| Modulation Of Chemical<br>Synaptic Transmission<br>(GO:0050804) | 7.51E-07 | 3.52 |
| Neuron Projection Development<br>(GO:0031175) | 7.78E-07 | 2.84 |
| Signal Release From Synapse<br>(GO:0099643) | 1.07E-06 | 7.79 |
| Regulation Of Neuron<br>Projection Development<br>(GO:0010975) | 1.07E-06 | 2.93 |
| Neuron Development<br>(GO:0048666) | 3.49E-06 | 3.02 |
| Axon Development<br>(GO:0061564) | 1.57E-05 | 3.51 |

(b) Representative Ontology Categories (RIN Negatively Correlating Genes)
| **Ontology Categories** | **FDR-adjusted**<br>**p-value** | **Odds Ratio** |
| --- | --- | --- |
| Response To Cytokine<br>(GO:0034097) | 9.02E-06 | 4.20 |
| Regulation Of I-kappaB<br>kinase/NF-kappaB Signaling<br>(GO:0043122) | 9.56E-05 | 2.86 |
| Defense Response To Symbiont<br>(GO:0140546) | 6.20E-04 | 3.23 |
| Positive Regulation Of I-kappaB<br>kinase/NF-kappaB Signaling<br>(GO:0043123) | 0.00125 | 2.86 |
| Defense Response To Virus<br>(GO:0051607) | 0.00201 | 2.70 |
| Positive Regulation Of<br>Cytokine-Mediated Signaling<br>Pathway (GO:0001961) | 0.0133 | 4.94 |
| Regulation Of NIK/NF-kappaB<br>Signaling (GO:1901222) | 0.0173 | 3.40 |
| Cellular Response To Cytokine<br>Stimulus (GO:0071345) | 0.0186 | 2.04 |
| Positive Regulation Of Defense<br>Response To Virus By Host<br>(GO:0002230) | 0.0186 | 6.23 |
| Cellular Response To<br>Mechanical Stimulus<br>(GO:0071260) | 0.0195 | 4.39 |

## Discussion

RNA quality in brain autopsy tissue may be influenced by a variety of factors, including peri-mortem conditions that influence pH (4, 6, 7, 12, 13). Here, we conduct a study across multiple datasets relating RIN and PMI to post-sequencing RNA-seq quality control metrics and find that PMI is minimally useful for predicting RNA quality using metrics derived from RNA-seq data.

The majority of the prior work that has been done using non-sequencing based analysis suggests that PMI has no to minimal value for predicting RNA quality (4, 6-9), although some publications suggest that PMI does matter to varying degrees (3, 10). Notably, Birdsill et al. found that 4 genes (ADAM9, LPL, PRKCG, and SERPINA3) negatively correlated with PMI in human brain tissue (out of 85 tested) (3), and Durrenberger et al. found that 18S rRNA negatively correlated with PMI (out of 7 reference genes tested) (11). At the individual gene level, we find that PMI positively correlates with 138 genes in common across the 3 autopsy datasets. While this is a small number in comparison to RIN, it is also interesting that all of these genes positively associate with PMI. While no ontology categories are significant, one could speculate that these genes may represent post-mortem changes in transcription associated with dying tissue, which is different from the RNA degradation that is normally assumed to the primary artifact associated with PMI.

Indeed, PMI also does not correlate with 3’ or 5’ bias, which is a common measurement of RNA degradation. This is particularly striking in the ROSMAP dataset with poly-A selection, where 3’ and 5’ bias correlate with RIN, but do not correlate with PMI. However, PMI does consistently correlate with PTT in two out of the three autopsy datasets, suggesting that reduced transcriptomic complexity relates to PMI in some fashion. We also note that PTT is a general barometer of RNA health, correlating with both RIN and 3’ and 5’ bias consistently. This makes PTT is the broadest QC metric we analyzed, as PTT captures variation in RNA quality that is observed in RIN, DV200, and measurements of 3’ and 5’ bias. PMI also significantly correlates with RIN in two of the three autopsy datasets, which suggests that RIN may be modulated by variables that in turn are not significantly correlating with many of the post-sequencing RNA quality metrics associated with RIN.

In contrast to PMI, RIN correlates with 3’ and 5’ bias, as well as with several thousand genes across all four datasets. Of particular interest is the fact that neuronal function ontology groups positively correlate with RIN, while immune response ontology groups negatively correlate with RIN. The fact that genes negatively correlate with RIN at all is itself counterintuitive, given that declining RIN scores and accompanying RNA fragmentation would be predicted to lead to loss of genes, not increases in genes that negatively correlate with these phenomena. Although speculative, one possible way to think about this is to not assume that RIN is driving these changes, but perhaps the other way around. As noted earlier, peri-mortem factors (for example sepsis) influence RNA quality (4, 6, 7, 12, 13). While this explanation obviously doesn’t apply to the NPH biopsies that also show these changes (as well as autopsy cases that didn’t die with sepsis), a more general phenomenon may be that brain inflammation, regardless of cause, contributes to lower RNA integrity. While it remains unclear exactly why inflammatory transcriptomic changes are elevated in brain tissue with lower RIN scores, the fact that these changes are found in common across four datasets suggests this is a reproducible phenomenon that is worth exploring in future work.

In summary, we have comprehensively assessed the impact of PMI and RIN on post-sequencing quality control metrics in brain tissue. Our results are overall consistent with prior work that has been done in this area with qPCR and microarray analysis, and give a broader assessment of the marginal but nevertheless reproducible impact of PMI on RNA quality. In addition, we also identify genes that are most likely to be impacted by RIN in RNA-seq studies. These findings will be of interest to researchers using human brain tissue for transcriptomic analysis, and also of potential interest to brain bank staff banking cases of interest with longer PMIs.

## Acknowledgements

This work was supported by NIH grants K08AG049938, K76AG054868, R01AG073360, R01AG072474, P30AG066462, P30AG10161, P30AG72975, R01AG15819, R01AG17917, U01AG46152, and U01AG61356, and with support from The Thompson Family Foundation.

